# Ultrasensitive proteomics uncovers nociceptor diversity and novel pain targets

**DOI:** 10.1101/2025.09.20.677124

**Authors:** Sampurna Chakrabarti, Anuar Makhmut, Atena Mohammadi, Wenhan Luo, Lin Wang, Gary R. Lewin, Fabian Coscia

## Abstract

The richness of our somatosensory experience is reflected in the functional diversity of somatic sensory neurons. Single-cell sequencing (scRNA) of sensory neurons has revealed a molecular basis for such diversity^1–3^. However, sensory neuron diversity has yet to be captured at the level of the proteome. Here, we combined electrophysiology with deep visual proteomics (DVP)^4^ to quantify over 6,000 proteins from phenotypically-defined sensory neurons and identified novel proteomic markers of sensory neuron subtypes. Comparative analysis revealed both concordance and meaningful divergence between transcriptomes and proteomes. We further show that up to 3,000 proteins can be quantified from one-fourth of a single neuron, demonstrating subset-specific protein signatures. In culture, nociceptive neurons can be acutely sensitized to mechanical stimuli by nerve growth factor (NGF) which normally drives inflammatory pain *in vivo*^5^. Indeed, overnight exposure of peptidergic nociceptors to NGF and a protein kinase C (PKC) activator produced functional sensitization associated with proteome changes. Functional knockdown experiments identified the up-regulated B3GNT2 enzyme as a potential effector of nociceptor sensitization. In summary, we present a high-resolution proteomic resource linking molecular identity to function, enabling the discovery of novel mechanisms underlying somatic sensation and pain sensitization.

## Introduction

Somatic sensation involves a rich diversity of perceptions, ranging from light touch, limb position, coolness, and warmth. However, sensations can also be acutely or chronically painful with distinct qualitative features depending on the precipitating insult, such as burns or traumatic injury. Such qualitatively different sensations are initiated by activation of sensory neurons with their cell bodies in the dorsal root ganglia (DRG). The DRG was the first neuronal tissue to be analyzed using single-cell RNA sequencing (scRNA seq) and the results revealed a rich diversity of sensory neuron sub-types that can be identified across species^1–3,6,7^. Such data have enabled a deeper understanding of the sensory neuron diversity that contributes to the richness of somatic sensation. However, while scRNA seq data can be used to describe cellular phenotypes and developmental trajectories^8^ it cannot, by itself, predict the cellular proteome which determines functional phenotypes^9,10^. Independent of transcription, multiple mechanisms exist to dynamically regulate the proteome, with timescales ranging from minutes to days^11,12^. Bulk proteomics has been used to characterize protein changes in sensory tissue involved in inflammatory and neuropathic pain^13–16^. However, such data cannot be directly mapped onto different cell types which are equipped with their own specific signaling modules that mediate functionally relevant plasticity. Cell specific signaling modules can only be accessed by obtaining high resolution proteomes of single cell types. Recent breakthroughs in sample processing, ultrasensitive mass spectrometry, and computational proteomics have now made this feasible ^4,17^.

Here, we used deep visual proteomics (DVP)^4^, which combines artificial intelligence (AI)-guided image analysis, automated laser microdissection (LMD), and ultrasensitive mass-spectrometry to obtain the proteomes of distinct sensory neuron cell types. We describe the first quantitative proteomic comparison of mouse sensory neuron subsets, quantifying 5-6 thousand unique proteins per cell type. Sensory neurons that detect noxious stimuli are termed nociceptors, and the activation and sensitization of these neurons is necessary for acute and chronic pain^18^. There is considerable functional diversity amongst nociceptors, and one classic example is the distinction between peptidergic and non-peptidergic nociceptors^19–21^. These two nociceptor subsets possess distinct functional properties^21,22^, developmental trajectories^23,24^ and even regenerative capacities^25,26^. Importantly, peptidergic nociceptors, but not non-peptidergic nociceptors, are sensitive to nerve growth factor (NGF), which is a key mediator of chronic inflammatory pain in animals and humans^27–29^. We first focused our approach on electrophysiologically characterized peptidergic and non-peptidergic mouse neurons in culture.

Deep proteomes for each cell type were obtained from 30-50 cells with high biological reproducibility across animals. We found high concordance between proteomes obtained from cultured neurons and the same neuronal cell bodies taken from DRG tissue sections. Using a cellular model of acute NGF and phorbol ester mediated sensitization, we show how only the peptidergic nociceptors display a rapid increase in their mechanosensitivity. This cellular model mimics nociceptor sensitization mediated by NGF *in vivo*, which has a rapid onset, but is also long-lasting^5^. We went on to identify dynamic protein changes associated with nociceptor sensitization and identified the glycosyltransferase enzyme B3GNT2, as being causally involved in nociceptor sensitization. Thus, our methodology was validated as a powerful new tool that complements and extends the existing sc-RNA seq datasets of neuronal diversity. Moreover, we show how deep neuronal subtype-specific proteomics can identify novel molecular drivers of pain hypersensitivity.

## Results

### Sensory neuron-subtype specific proteomes

Many, but not all cultured sensory neurons possess mechanically-gated currents^22,30–32^. Patch clamp electrophysiology shows that almost all large diameter mouse sensory neurons (>30 µm), which have an electrophysiological signature of mechanoreceptors, possess mechanically-gated currents to indentation stimuli^30–33^. In contrast, small-medium diameter (<30 µm) sensory neurons with a nociceptor-like electrophysiological signature exhibit mechanically gated currents with slower kinetics and reduced frequency compared to mechanoreceptors^30,32,34^. Peptidergic and non-peptidergic nociceptors exhibit distinctive biophysical properties, including responsiveness to mechanical stimuli^21,22^. We used isolectin-B4 coupled to AlexaFluor-594, which binds to α-D-galactose carbohydrate residues on the surface of non-peptidergic nociceptors, to live label these cells in culture^21,35^ (**Fig 1a,b**). The incidence of mechanically gated currents evoked by cell membrane indentation was significantly lower in IB4^+^ nociceptors (10%) than in IB4^-^ nociceptors (∼50%) (IB4^-^, 12/23 vs. IB4^+^, 2/18, p = 0.005, chi-sq test **Fig 1c, d, Extended Data Fig 1a**). Approximately half of the mechanically evoked currents in nociceptors were rapidly adapting as has been described before^32,34^ (Extended Data Fig 1b). IB4^+^ nociceptors also showed a tendency to have broader action potentials (half-peak duration p = 0.14, unpaired t-test with Welch’s correction) with a larger (p = 0.04, unpaired t-test) afterhyperpolarization peak amplitude (**Fig 1e**). The two IB4^+^ neurons with a mechanosensitive current had action potential properties typical for this neurochemical sub-type (Fig 1e). Therefore, IB4^+^ non-peptidergic nociceptors are functionally distinct from IB4^-^ peptidergic nociceptors in mice, as previously observed in rats ^21,35^.

**Fig 1:**
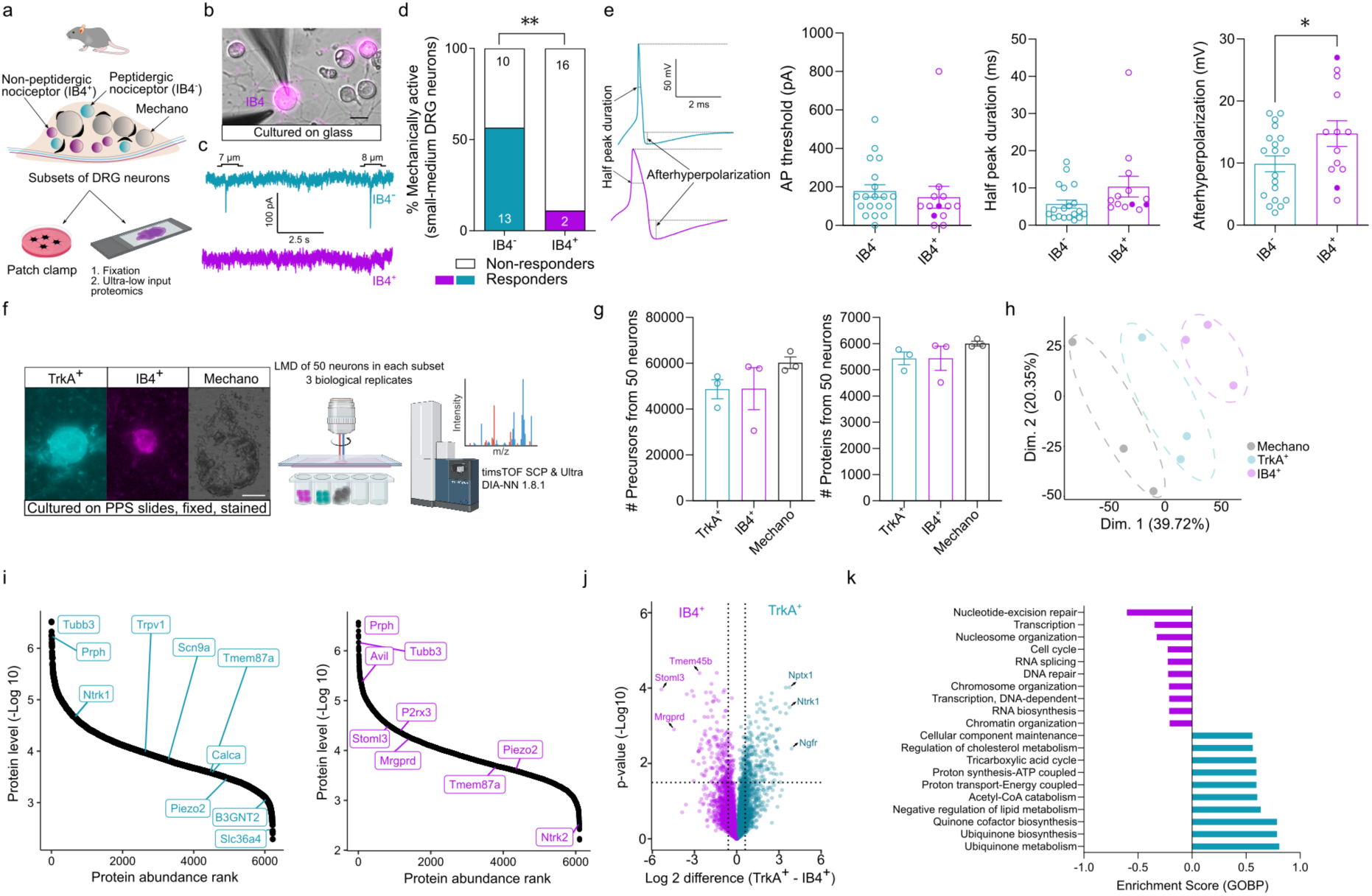
Electrophysiology-guided, subset-specific proteome of cultured sensory neurons. **a)** Schematic workflow showing patch clamp electrophysiology. **b)** IB4 labelling in magenta, (scale = 20 µm) and ultra-low input proteomics of cultured sensory neurons. **c)** Representative traces of indentation-evoked, mechanically-activated currents in an IB4^-^ peptidergic nociceptor and the absence of such currents in an IB4^+^ non-peptidergic nociceptor. **d)** Percentage of mechanically sensitive IB4^-^ and IB4^+^ neurons. The number of neurons is indicated in the bars. ** = p < 0.01, chi-sq test. **e)** Representative action potential (AP) traces recorded from an IB4^-^ and an IB4^+^ nociceptor, with corresponding quantification of AP threshold, half peak duration, and afterhyperpolarization of the measured APs. Each dot represents a single neuron. Filled dots in IB4^+^ group represents the two mechanosensitive non-peptidergic neurons. Data were obtained from both male and female mice. * p < 0.05, unpaired two-tailed Student’s t-test. **f)** Representative images of sensory neurons cultured and stained on laser microdissection (LMD)-compatible membrane slides prior to LMD and mass spectrometry analysis (scale = 20 µm). **g)** Numbers of precursors and proteins identified from peptidergic (IB4^-^/TrkA^+^) and non-peptidergic (IB4^+^/TrkA^-^) nociceptors and mechanoreceptors. Each dot represents a male mouse. **h)** Principal component analysis of peptidergic and non-peptidergic nociceptors and mechanoreceptors. **i)** Dynamic range of protein abundance in the two nociceptor subsets. **j)** Volcano plot showing pairwise proteomic comparison between nociceptor subsets with marker proteins highlighted. **k)** Pathway enrichment analysis based on t-test differences between nociceptor subsets. The top ten pathways with a Benjamini-Hochberg false discovery rate (FDR) < 0.05 are shown. Cyan = peptidergic, magenta = non-peptidergic, black = mechanoreceptor. Error bars represent s.e.m.

To profile the global proteomes of these functionally distinct nociceptor subsets, we utilized our recently developed ultra-low-input proteomics workflow, integrating imaging and laser microdissection (LMD)^17^. A portion of the mouse sensory neurons assessed with electrophysiology was also cultured overnight on LMD-compatible slides, fixed, and immunolabeled with antibodies against the cell surface tyrosine kinase receptor, TrkA, and IB4, two markers that largely discriminate between these two nociceptor populations, peptidergic (TrkA^+^, IB4^-^) and non-peptidergic (IB4^+^, TrkA^-^)^23^ (Fig 1f). From each of the three mice, we collected 50 TrkA^+^ and 50 IB4^+^ neurons using LMD. We additionally included 30 large diameter mechanoreceptors that do not normally stain positive for nociceptor markers, such as IB4 and TrkA. Proteomic measurements were conducted using a trapped ion mobility mass spectrometer (Bruker timsTOF SCP/Ultra) operated in data-independent (diaPASEF) acquisition mode. We identified more than 40,000 precursors and 6,000 proteins across all three subsets of sensory neurons, with biological replicates closely grouping together in principal component analyses (**Fig 1g, h**). Among the quantified proteins were ∼ 400 transmembrane proteins, including many ion channels associated with nociception, such as Trpv1, Trpa1, and Scn9a (**Supplementary Table 1**). Hierarchical clustering of all 821 ANOVA-significant proteins revealed clearly demarcated clusters with unique protein signatures among the three subsets, enabling us to define the first proteomics-based signature of peptidergic and non-peptidergic nociceptors, as well as mechanoreceptors (**Extended Data Fig 1c, Supplementary Table 2**). Interestingly, among the top markers (**Extended Data Table 1**) were proteins such as Hpcal1 (a neuron-specific, calcium binding protein) and Armcx1 (a mitochondrial protein), which we found to be novel subtype-specific markers for peptidergic and non-peptidergic neurons, respectively.

The most abundant proteins quantified were Tubb3, Prph, and classical markers of sensory neurons^36^, implying minimal admixing with non-neuronal cell types, such as satellite cells^37^ (**Fig 1i**). Furthermore, many well-known markers of the three different subsets were quantified and distributed over a dynamic range spanning more than four orders of magnitude (highlighted in **Fig 1i and Extended Data Fig 1d**). The proteome of IB4^+^ neurons differed significantly from that of TrkA^+^ neurons, with the largest differences in proteins encoded by well-established marker genes^3^ , such as *Mrgprd* (in IB4+ population), *Ntrk1* (in TRKA^+^ population), and *Nefh/Ntrk3* (in mechanoreceptors) (**Fig 1j and Extended Data Fig 1e**). Notably, Tmem45b^38^, a recently identified membrane protein implicated in mechanical pain hypersensitivity in the non-peptidergic population, was one of the most highly expressed proteins in IB4^+^ neurons in our dataset. Our subset-specific proteome analysis also revealed differential protein abundances between peptidergic and non-peptidergic clusters for genes with very low mRNA abundance, such as Stoml3 (a regulator of sensory mechanotransduction^39^) and the synaptic remodeling protein, Nptx1^40^. Pathway enrichment analysis identified terms associated with “metabolism” and “biosynthesis” processes as significantly associated with the TrkA^+^ neuronal proteome (FDR < 0.05), potentially reflecting their superior capacity to sprout centrally in the dorsal horn^25^ (**Fig 1k**). Similar gene ontology-based enrichment analysis between IB4^+^ nociceptors and mechanoreceptors revealed “C-fiber” as the most significantly enriched term in the IB4^+^ neuronal proteome, validating the premise of such an analysis (**Extended Data Fig 1f**). In conclusion, our approach shows that it is possible to uncover deep proteomes from sensory neuron subtypes, a rich resource for the discovery of nociceptor-specific pain targets.

### Common proteomic signatures of intact and cultured neurons

Cultured sensory neurons are amenable to a host of electrophysiological, biochemical, and functional manipulations and are an important model for pain research^41^. However, questions remain on how the culturing process changes gene expression and protein levels when compared to intact DRG neurons. A comparison between cultured and intact DRGs using bulk RNA sequencing revealed an overall similarity between these conditions, but such an analysis at the protein level is missing^42^. To address this, we placed 5µm thick cryosections from intact mouse lumbar DRGs onto LMD-compatible membrane slides and immunostained them for calcitonin gene-related peptide (CGRP) and P2X purinoceptor 3 (P2X3) (**Fig 2a**). These two markers, which largely overlap with TrkA^+^ and IB4^+^ cells, respectively, show little species-specific variability and are good markers for peptidergic and non-peptidergic neurons^43–45^. To improve the efficiency of sample throughput, we used a custom-trained deep learning-based Cellpose 2.0^46^ model to segment CGRP^+^ and P2X3^+^ neurons. The generated segmentation masks were dilated radially by 10 µm using the image analysis software Qupath^47^ to ensure that the entire neuron profile was included. These contours were manually inspected to ensure no overlap between adjacent neurons, followed by automated LMD, as per our deep visual proteomics (DVP) workflow^17^. Neurons double positive for CGRP and P2X3 were excluded. We collected 100 CGRP^+^ and 100 P2X3^+^ nociceptor sections pooled from each of the three mice to quantify ∼25000 precursors and ∼3500 proteins from each subset (**Extended Data Fig 2a, Supplementary Table 3**). A similar proteome depth was obtained from large-diameter mechanoreceptors in a separate experiment (**Extended Data Fig 2b, c, Supplementary Table 4**). Importantly, we found that over 90% of the proteins quantified in both peptidergic and non-peptidergic groups in intact sensory neurons were also detected in cultured sensory neurons (**Fig 2b**). The lower depth of proteome compared to cultured neurons is likely due to the lower volume of samples collected from tissues. A plot of the differentially regulated proteins in peptidergic vs. non-peptidergic neurons from tissue (CGRP^+^ vs. P2X3^+^) and cell culture (TRKA^+^ vs. IB4^+^) showed excellent correlation with *Calca* (gene name for CGRP) being the top regulated protein in the peptidergic cluster (Pearson correlation = 0.73, **Fig 2c**). This finding confirms that the proteomic identity of peptidergic and non-peptidergic neurons remains consistent, regardless of the markers used for classification. This also suggests that the culturing procedure had only a minimal influence on the proteomic identity of the neurons. Gene ontology (GO) enrichment analysis of the 315 and 261 unique proteins in the CGRP^+^ and P2X3^+^ subsets of sensory neurons showed “extracellular matrix structure” as a highly significant term, which is consistent with intact sensory neurons being embedded in their native matrix (**Fig 2d**). In contrast, the GO terms associated with the unique proteins identified from cultured nociceptors (Extended Data Fig 2d) were “membrane coat” as well as metabolism related terms in the peptidergic subset similar to Fig 1k. “Extracellular matrix” was also amongst the most significant enrichment term for the 650 unique proteins in the intact NF200^+^ mechanoreceptors (**Extended Data Fig 2e**).

**Fig 2:**
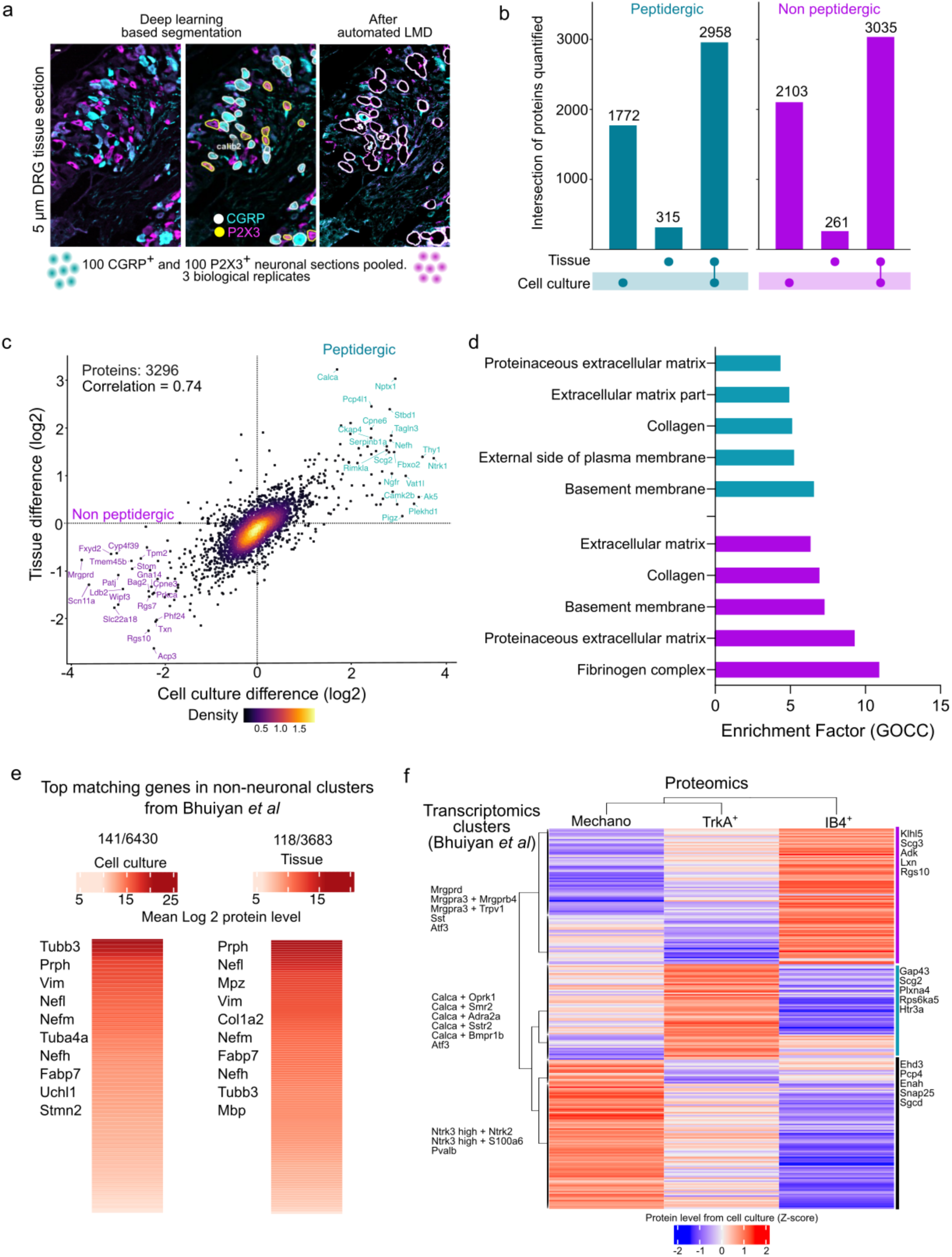
Subset-specific intact nociceptor proteomics from tissue sections. **a)** Representative images of a 5 µm intact dorsal root ganglion (DRG) tissue section stained with CGRP and P2X3, segmented using a Cellpose 2.0 model and processed by semi-supervised, automated LMD. “calib2” denotes one of the calibration points used to transfer masks from Cellpose to LMD. **b)** Upset plot showing common and unique proteins quantified from nociceptor subsets in intact tissue and culture. **c)** Correlation of differences in protein levels between nociceptor subsets for common proteins identified in both cultured and intact sensory neurons, along with a color gradient of the relative point density. **d)** Pathway enrichment analysis (Fischer’s exact test) of uniquely identified proteins from intact tissue-derived nociceptors. The top five pathways with a Benjamini-Hochberg false discovery rate (FDR) < 0.05 are shown. **e)** Heat map of protein levels quantified in cultured and intact sensory neurons, corresponding to genes identified in non-neuronal clusters in the harmonized single-cell transcriptomics atlas (Bhuiyan et al.^3^). Each row represents the mean Log2 protein level across all three sensory neuron subsets of a given matched gene. **f)** Heat map showing overlap between the combined top 50 marker genes from all subsets of the harmonized single-cell transcriptomics atlas and our subset-specific proteome from cultured neurons. Each row represents the z-score of the protein level of a matched gene. Cyan = peptidergic, magenta = non-peptidergic.

Next, we compared our subset-specific proteomes to existing scRNA seq-based sensory neuronal clusters. For this analysis, we utilized a recently published database from Bhuiyan and colleagues^3^, which harmonized existing sensory neuron and associated non-neuronal single-cell databases comprising both single-cell and single-nucleus RNA-seq data (referred to hereon as scRNA seq data). Among the 1556 unique genes quantified in non-neuronal cells via scRNA seq, only ∼150 genes matched our proteome dataset. Therefore, we estimated that ∼ 3% of our measured proteomes may have been derived from non-neuronal cells, such as satellite and Schwann cells, which are tightly attached to neurons in sections or in culture (**Fig 2e**). Notably, genes like Tubb3 and Prph are detected in both neurons and Schwann cells and therefore 3% is probably an overestimation of the non-neuronal contribution.

Next, a compiled list of the top 50 marker genes from the 18 sensory neuronal subclusters in the harmonized scRNA seq database was matched to our deep proteome from cultured neurons. Among the 554 unique genes present in the transcriptomic marker list, 255 matched our proteomes, revealing well matching clusters for cell type-specific transcriptomes and proteomes (**Fig 2f**). For example, transcriptomics based peptidergic markers, *Gap43 and Scg2*, were also highly expressed at the protein level in our TrkA^+^ peptidergic group. Taken together, our ultra-low input, subset specific proteome closely aligns with the sc-RNA seq-derived transcriptomic signatures of sensory neuron subsets.

We additionally compared the transcript/protein detectability between the ∼15000 gene deep scRNA seq dataset of Sharma and colleagues^2^ and our ∼6000 protein deep cultured neuron data. Proteins with higher transcript counts in the scRNAseq study were more likely to be detected by proteomics (Extended Data Fig 3a). For example, we detected 80% of genes with a transcript read count 30 or higher, which compared to 60% of the genes with ∼5 average transcript reads. However, even genes with low transcript levels (e.g., Mrgprd) could be identified by proteomics as nociceptor subset specific markers (Extended Data Fig 3b).

### Single neuron proteomes delineate nociceptor subsets

Next, we investigated whether our method is capable of quantifying single intact sensory neuron proteomes and whether these data capture subset specificity. We laser microdissected single CGRP^+^ and P2X3^+^ nociceptor sections from DRGs of each of the three mice (**Fig 3a**). After quality control, 20 CGRP^+^ peptidergic neurons and 22 P2X3^+^ non-peptidergic single neurons remained in the dataset, with peptidergic and non-peptidergic nociceptors forming distinct clusters (**Fig 3b**). The highest number of proteins quantified from a single nociceptor was 1691, from an average of ∼3000 precursors (**Extended Data Fig 4a**). A comparison of differentially regulated proteins between peptidergic and non-peptidergic nociceptors from 100 pooled contours *versus* a single contour showed a high global proteome correlation, with CGRP (*Calca)* being one of the highest regulated proteins in the peptidergic subset (Pearson correlation = 0.6, **Fig 3c**). As expected, the quantified proteins from single neuron slices were the most abundant proteins and were part of the stable neuronal core proteome^48^. We demonstrated the existence of such a stable core proteome in a single section of a neuron, while preserving subset specificity. Comparing peptidergic and non-peptidergic single neurons revealed distinct differentially abundant proteins, with 32 significantly upregulated proteins in peptidergic neurons and 25 in non-peptidergic neurons (**Fig 3d, Supplementary Table 5**). By searching the open-source drug-gene interaction database (DGIdb, v.5.0.8) against upregulated proteins in the single peptidergic neuron proteome, we found interactions between *Hspb1* and two approved analgesics: the non-steroidal anti-inflammatory drug (NSAID) rofecoxib and argipressin. Additionally, *Eno2*, which is upregulated in the peptidergic cluster, interacts with gabapentin, a widely used analgesic, and indomethacin, an NSAID. Similarly, among the differentially upregulated proteins in the non-peptidergic cluster, *Clgn* interacts with the anti-rheumatic agent leflunomide. Gene ontology analysis further showed that the proteome of single CGRP^+^ peptidergic nociceptors were enriched in “metabolism” -related terms like those observed with our deep proteome from 50 cultured nociceptors, while “glycosylation” was associated with non-peptidergic neurons in agreement with their IB4 (lectin) binding capacity (**Fig 3e**). Previous scRNA seq studies identified six subclusters of CGRP^+^ neurons, prompting us to explore whether we could probabilistically map single CGRP^+^ neuronal proteomes to these clusters. Based on the Euclidean distance between normalized positive fold change values from transcriptomics and protein levels of the matched marker genes, we found that our pooled 50 cultured peptidergic nociceptor proteome best matched the “Calca+Dcn” transcriptome cluster, with undefined function^3^ (**Extended Data Fig 4b**). A similar comparison of single CGRP^+^ neurons with transcriptomic clusters revealed that some neurons also had a probability of belonging to other clusters such as “*Calca+Adra2a*” or “*Calca+Smr2*”. Single neurons showed the lowest likelihood of belonging to the rare cell type “*Calca+Oprk1*” or Aδ-nociceptors “*Calca+Bmpr1b*”. Therefore, our analysis suggests that single neuron proteomes can, in principle, be used to generate unbiased clustering of sensory neurons and their function can be probed directly using electrophysiology to determine molecular determinants of function at a single neuron level.

**Fig 3:**
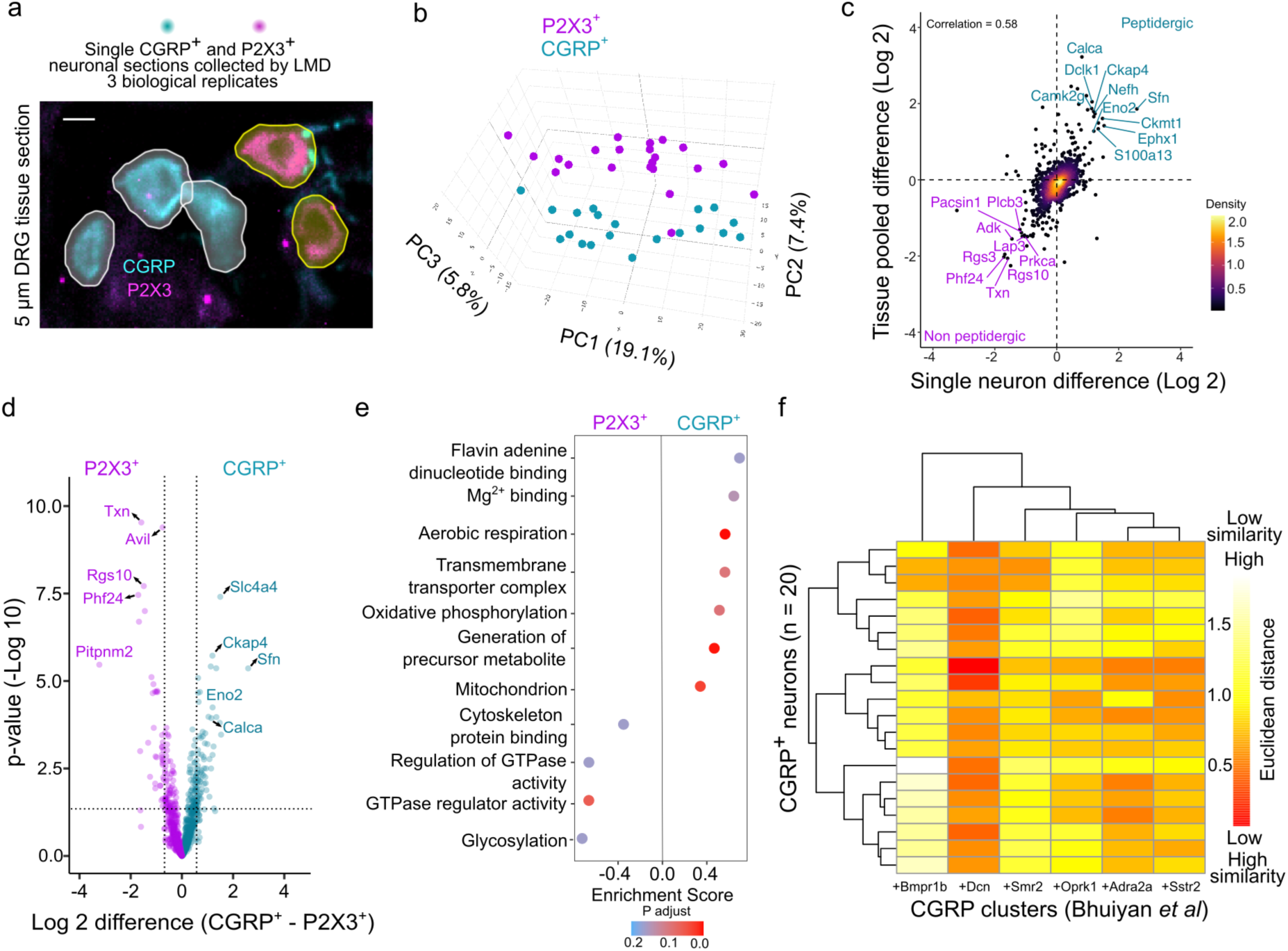
Proteome from a section of a single nociceptor. **a)** Representative image of a dorsal root ganglion (DRG) section stained with CGRP (peptidergic) and P2X3 (non-peptidergic) and the corresponding Cellpose derived segmentation masks. **b)** A 3D principal component analysis of individual CGRP^+^ and P2X3^+^ nociceptors. **c)** Correlation of differences in protein abundance between nociceptor subsets for common proteins identified in both pooled and single sensory neuron sections, along with a color gradient of the relative point density. **d)** Volcano plot showing pairwise proteomic comparison between combined single nociceptor proteomes, with marker proteins of each subset highlighted. **e)** Pathway enrichment analysis based on t-test differences between nociceptor subsets. The top pathways enriched in the peptidergic and non-peptidergic neurons are shown. **f)** Similarity plot based on Euclidean distance between enriched genes in CGRP (peptidergic) subclusters (columns) identified by single-cell transcriptomics and proteomics of individual CGRP^+^ nociceptors (rows). Red = high similarity; white = low similarity. Cyan = peptidergic, magenta = non-peptidergic.

Proteome coverage was significantly higher in large diameter mechanoreceptors (n = 36), with a range of 1,404-3,615 identified proteins (**Extended Data Fig 4c, Supplementary Table 3**). We expect that advances in sample preparation techniques and higher-sensitivity MS will further enhance proteome depth from single neurons. Our analysis shows that, at present, a pooled approach of 50 phenotype matched cultured neurons yields the best proteomic depth, which may be leveraged to identify novel pain targets.

### A cellular model of mechanical pain hypersensitivity

NGF is an important mediator of painful mechanical hypersensitivity in humans and rodents and acts by downstream signaling via its receptor TrkA^5^. We next hypothesized that our neuronal subtype-resolved proteomics approach could identify downstream proteins involved in NGF-dependent mechanical hypersensitivity. We used an *in vitro* “inflammation” model in which isolated nociceptors were exposed to nerve growth factor (NGF) and the PKC activator phorbol 12-myristate 13-acetate (PMA) (**Fig 4a**). As previously reported in rat sensory neurons^22^, overnight treatment with 100 ng/ml NGF and 10 nM PMA doubled the proportion of neurons with indentation-induced currents (control, 12/23 vs. NGF-PMA, 19/22, p=0.01, chi-sq test, **Fig 4b**). A non-significant trend towards increased peak current amplitude was also observed when compared to the control (p = 0.2, unpaired t-test with Welch’s correction, **Fig 4b, Extended Data Fig 5a**). The proportion of rapidly, intermediately and slow-adapting currents were similar in control and sensitized states (**Extended Data Fig 5a**). Notably, the proportion of neurons in which residual non-adapting inward current was observed after removal of indentation increased from 16.6 % in control to 52.6 % in inflamed neurons (p = 0.04, chi-sq test, **Fig 4c**). No changes in the action potential firing threshold or threshold for mechanically activated currents were observed (**Extended Data Fig 5b**). The sensitizing effects of NGF and PMA were restricted to small-medium diameter IB4^-^, peptidergic neurons, consistent with their expression of the TrkA receptor (**Fig 4d, Extended Data Fig 5c**). Therefore, we hypothesized that our phenotype-resolved discovery proteomics pipeline of inflamed versus control TrKA^+^ nociceptors might reveal regulated proteins with a causal role in nociceptor sensitization.

**Fig 4:**
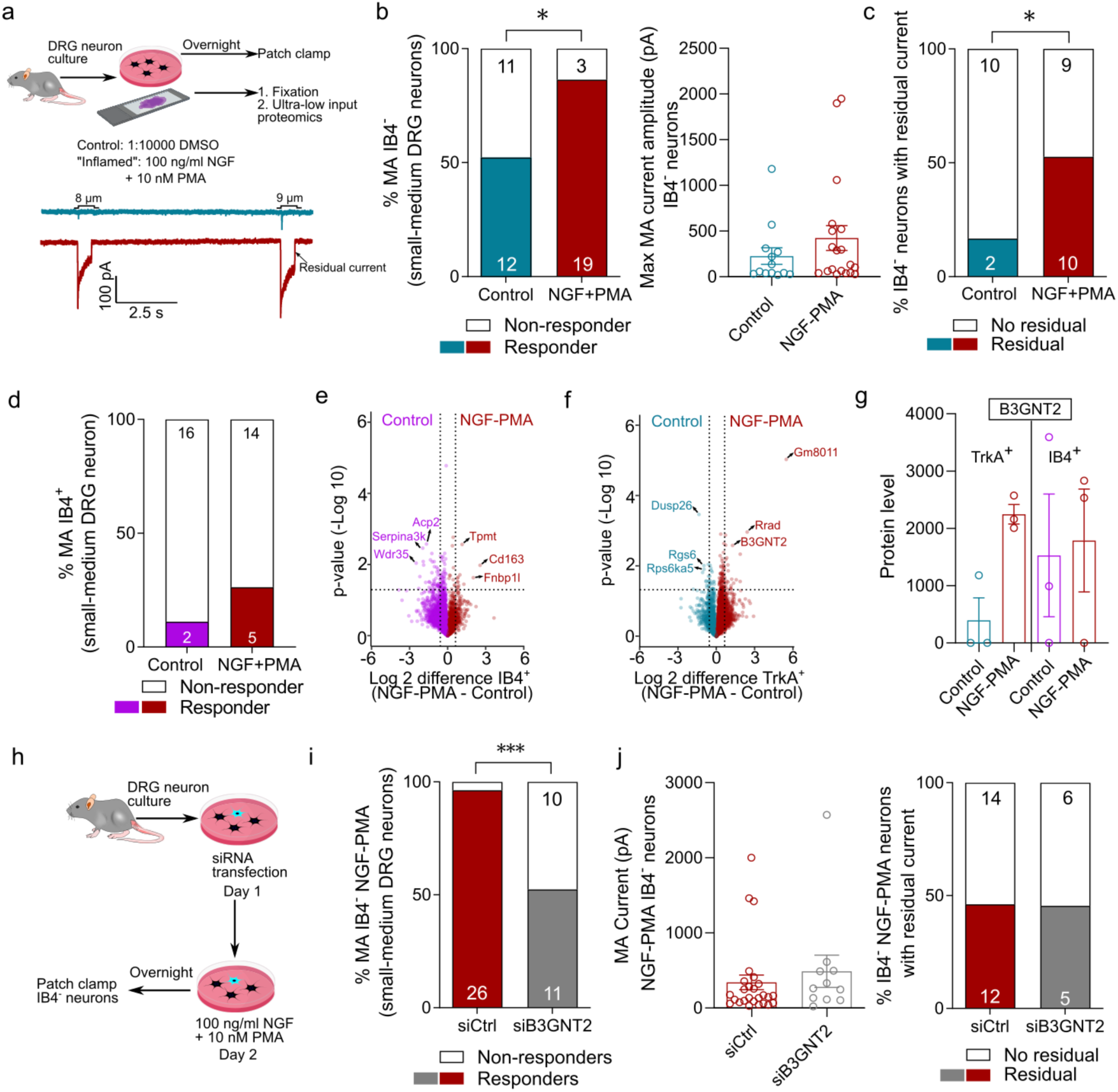
B3GNT2 is identified as a driver of peptidergic neuron sensitization. **a)** Schematic workflow of sensory neurons cultured in an “inflammatory soup” containing nerve growth factor, NGF, and phorbol 12-myristate 13-acetate, PMA (top), along with representative indentation-evoked currents from a control (cyan) and an inflamed (red) peptidergic neuron. **b)** Percentage of mechanically sensitive control and inflamed peptidergic nociceptors (left). The number of neurons is indicated in the bars, along with their corresponding current amplitude (right). **c)** Percentage of control and inflamed peptidergic nociceptors with residual current. **d)** Percentage of mechanically sensitive control (magenta) and inflamed (red) non-peptidergic nociceptors. The number of neurons is indicated in the bars. * = p < 0.05, chi-sq test. Volcano plots showing the pairwise proteomic comparison between control and inflamed neurons in non-peptidergic **(e)** and peptidergic **(f)** nociceptors with top regulated proteins highlighted. **g)** Bar plots showing raw protein levels of B3GNT2 in the control and inflamed groups of peptidergic (TrkA^+^) and non-peptidergic (IB4^+^) nociceptors. Each dot represents a single mouse. **h)** Schematic workflow of the validation experiment by knockdown of *B3gnt2* using siRNA prior to inflammation and patch-clamp electrophysiology of peptidergic neurons. **i)** Percentage of mechanically sensitive inflamed peptidergic nociceptors transfected with control siRNA (red) and B3gnt2-targeting siRNA (grey). The number of neurons is indicated in the bars. *** = p < 0.001, chi-sq test. **j)** Current amplitudes (left) and percentage of neurons with residual currents in inflamed peptidergic nociceptors transfected with control siRNA (red) and B3gnt2-targeting siRNA (grey). Error bars represent s.e.m.

### The enzyme B3GNT2 is a putative effector of nociceptor sensitization

Differential analysis of NGF-PMA-treated versus control neurons identified nine upregulated proteins (**Fig 4e**) in IB4^+^, seven in Mechano **(Extended Data Fig 5d)** and 18 in TrkA^+^ neurons (**Fig 4f**) (**Supplementary Table 6**). All upregulated proteins were unique to each of the three sensory neuron subsets. We focused on the most significantly upregulated proteins in TrkA^+^ peptidergic nociceptors. The sequence information available for the candidate protein with the highest -Log10 p value, *Gm8011* (predicted gene locus in Chromosome 14), appeared to be incomplete in NCBI, and it was thus not selected for functional validation. The second most significant candidate was Rrad, a Ras-related glycolysis inhibitor and a calcium channel regulator. Recent studies have suggested that Rrad increases cell surface tension, which could enhance the activity of mechanosensitive channels, such as Piezo1 and Trek1^49,50^. To evaluate the role of Rrad in inflammation-induced mechanical hypersensitivity in peptidergic neurons, we used siRNA directed against *Rrad* prior to nociceptor sensitization with NGF and PMA treatment (**Extended Data Fig 5e**). Non-targeting siRNA was used as a control and here the effects of NGF and PMA on mechanically gated currents were similar to controls with 25/26 “inflamed” neurons showing whole cell currents to cell indentation (**Fig 4a,b,c, Extended Data Fig 5f**). However, this NGF-PMA-induced mechanical hypersensitivity was not decreased in sensitized neurons after *Rrad* knockdown (**Extended Data Fig 5f,g**). This could be explained by a comparatively small number of sensory neurons that express Rrad, as assessed from scRNA seq studies^2,3^.

Our second candidate, B3GNT2^51^, an enzyme involved in the synthesis of poly-N-acetyllactosamine, was the third most upregulated protein in peptidergic-sensitized neurons. B3GNT2 was consistently detected in all three biological replicates of TrkA^+^ sensitized neurons, showing a two-fold increase in protein abundance compared to TrkA^+^ control neurons (only detected in one of the three samples) (**Fig 4g**). We transfected sensory neurons with siRNA targeting *B3gnt2,* followed by NGF-PMA treatment (**Extended Data Fig 5e, Fig 4h**). Compared to sensitized IB4^-^ peptidergic neurons transfected with control siRNA, *B3gnt2* knockdown reduced the proportion of neurons with mechanically activated currents to levels seen in untreated control nociceptors (11/21 responded to mechanical indentation, p = 0.0003, Chi-sq test, **Fig 4i**). No difference was found between control siRNA transfected and *B3gnt2* siRNA transfected IB4^-^ treated neurons in peak mechanically activated current amplitude or in the proportion of neurons with slowly inactivating residual currents, suggesting the role of other unknown proteins in NGF-induced mechanical hypersensitivity (**Fig 4j**). Nevertheless, our data identify a specific role for the B3GNT2 enzyme in the appearance of newly mechanosensitive peptidergic nociceptors following inflammatory stimuli. We did not investigate the role of B3gnt2 in IB4^+^ non-peptidergic neurons, since the protein was not differentially enriched in NGF-PMA treated IB4^+^ neurons. Our findings provide *in vitro* proof-of-concept validation for using ultra-low input proteomics to identify cell-type-specific proteomic changes directly relevant to pain physiology.

## Discussion

It has long been known that transcriptional signatures can significantly diverge from the proteome signature^10,12^. Here, we demonstrate how deep visual proteomics can be leveraged to obtain sensory neuron subtype-specific proteomes with depths equivalent to that of previous whole tissue proteomes^14^. This powerful technique allowed us to align cell-type specific proteomes to sensory neuron scRNA seq data.

Cultured sensory neurons are a robust model for investigating nociceptive mechanisms and screening for drug targets due to their relative ease of culture and functional manipulation^41^. However, concerns remain regarding whether culturing alters their fundamental cell state. While bulk sequencing data demonstrates minimal changes due to culturing^52^, these effects have not been thoroughly investigated at the protein level or in a neuronal subtype-specific manner. In the present study we demonstrate that the proteome of peptidergic and non-peptidergic nociceptors, as well as mechanoreceptors derived from cultured or intact DRG sections exhibit considerable overlap. The small number of unique proteins present in tissue sections are mostly derived from the extracellular matrix since these neurons are still embedded in their native matrix. Therefore, intact DRG neurons are preferable for studying neuronal interactions with endogenous extracellular matrix, while subset specific mechanisms can be investigated in both cultured and intact neurons.

We found that sensory neuron identity is ingrained in their proteomes, which can be defined by our ∼2,000 protein deep single neuron dataset obtained from tissue sections. This proteome coverage was comparable to that of cultured single-neuron proteomes^53^. The purpose of our single neuron proteome data was to demonstrate that, in principle, single-cell proteomics can complement sc-RNA seq classifications. We also demonstrate that obtaining meaningful electrophysiological and proteome data from the same sensory neuron is feasible, with future applications in matching proteomes to single cell physiology. The high sensitivity of our system also implies that quantifying proteins from the fine nerve endings is a future possibility.

Our ∼6400-protein dataset for three subtypes of sensory neurons was largely consistent with transcriptionally defined gene sets from the harmonized single-cell transcriptomics atlas^3^. Importantly, we define novel proteome-based markers of sensory neuron subsets, which complement single-cell transcriptomics derived markers. For example, only six genes (*Htr3a, Calca, Ntrk1, Atp2b4, Fgf12,* and *Dclk1*) in our top 50 proteome-defined peptidergic markers matched a compiled list of all top 50 genes across six Calca-expressing transcriptomic clusters. This highlights the importance of measuring cell type-specific proteomes that are not predictable from transcriptomes.

Here, we show how peptidergic and non-peptidergic nociceptive neurons have highly distinctive proteome signatures. Our data shed light on how the molecular make-up of these distinct nociceptor populations may determine their distinct anatomical connectivity and physiological properties^21–23,54^. The rapid appearance of mechanosensitivity in nociceptors that were previously unresponsive to mechanical stimuli is termed “unsilencing” and can be initiated by inflammatory mediators, such as NGF^5,55–57^. Nociceptor “unsilencing” is long known to be rapid in onset and long-lasting in a wide variety of nociceptors^58^, making it likely that changes in protein abundance or localization underlie this phenomenon. To illustrate the utility of our proteomics approach we asked whether agents that produce nociceptor “unsilencing” change protein abundance in defined sensory neuron sub-populations (**Fig. 4**). We showed that NGF combined with activation of protein kinase C (PKC) produced sub-type-specific nociceptor “unsilencing” in vitro^22^ (**Fig. 4**). Peptidergic sensory neurons, which predominantly express the NGF receptor *Ntrk1* (TRKA), were specifically sensitized to mechanical stimuli^22^. Following functional phenotyping, we identified significant changes in the proteome of stimulated peptidergic neurons that could, in principle, be effectors of inflammatory sensitization. We found that the beta-1,3-N-acetylglucosaminyltransferase enzyme B3GNT2^59^ was one such novel upregulated protein in sensitized nociceptors^51^. Using siRNA-mediated gene knockdown of *B3gnt2,* we could demonstrate reversal of inflammation-induced “unsilencing” of peptidergic neurons. Although the role of B3gnt2 in inflammatory acute and chronic pain requires further *in vivo* validation, our data implicate sugar modification of proteins as a necessary step in the acute mechanical sensitization of nociceptors. The loss of B3GNT2 activity in the mouse has been associated with mislocalization of axonal guidance factors in the olfactory system^60^, therefore, we speculate that the activity of this enzyme is necessary for the transport of mechanically gated ion channels or their regulators to the plasma membrane to mediate “unsilencing” in nociceptors. Interestingly, genetic variation at the *B3GNT2* locus in humans has been associated with more severe symptoms in patients with ankylosing spondylitis (a painful inflammatory arthropathy^61^). In contrast, knockdown of another candidate protein upregulated in sensitized nociceptors, Rrad, did not reverse inflammation-induced mechanical hypersensitivity, highlighting the importance of combining electrophysiology with ultra-sensitive proteomics in validation studies.

In summary, we provide a high-resolution, subset-specific proteome of functionally distinct sensory neurons. Importantly, we show that understanding the molecular pathways involved in nociceptor sensitization requires cell type-specific proteomes. Our dataset offers new opportunities to explore the molecular mechanisms underlying nociception, further advancing our understanding of how chronic pain develops and can be prevented.

## Materials and Methods

### Animals

Wildtype C57BL/6N adult (10-30 weeks) mice of both sexes were used in the study for electrophysiology and only male in proteomics experiments. N = 4 male and n = 3 female mice were used in patch clamp experiments in Figure 1. N = 3 male mice were used for both electrophysiology and cultured neuron proteomics. N = 2 female and n = 4 male mice were used for siCtrl patch clamp experiments. siB3gnt2 and siRrad patch clamp data was derived from n = 2 male and n = 1 female mice in each condition. Mice were housed in groups of 5 with food, water, and enrichment available ad libitum at the animal house of the Max Delbrück Center for Molecular Medicine (Berlin, Germany). All animal protocols were approved by the German federal authorities (State of Berlin).

### Dorsal root ganglion (DRG) neuron culture

Lumbar DRGs (L2-L5) were collected from mice into plating medium (DMEM-F12 (Gibco) supplemented with 10% fetal horse serum (FHS, Life Technologies) and 1% penicillin and streptomycin (P/S, Sigma-Aldrich)). These DRGs then underwent enzymatic digestion in 1.25% Collagenase IV (1 mg/ml, Sigma-Aldrich) for 1 hour at 37°C and 2.5% Trypsin (Sigma-Aldrich) for 15 min at 37 °C, followed by mechanical trituration with a P1000 pipette tip and purification in a 15% fraction V BSA column. The neurons were plated on glass-bottomed plates (for indentation assays) coated with poly-L-lysine and laminin for electrophysiology and in UV-sterilized (30 min) polyphenylene sulfide (PPS, Leica microsystems) membrane slides for neuroproteomics. Neurons were cultured at 37 °C in a 5% CO2 incubator.

For experiments involving in vitro inflammation, neurons were incubated overnight in 100 ng/ml nerve growth factor (NGF-ϕ3 human, N1408, Sigma, 0.1 mg/ml stock in water) and 10 nM phorbol 12-myristate 13-acetate (PMA, P1585, Sigma, diluted from a 1mM stock in DMSO) before whole cell electrophysiology.

### Patch clamp electrophysiology

Whole-cell patch clamp electrophysiology was performed using heat-polished borosilicate glass pipettes (Harvard apparatus, 1.17 mm x 0.87 mm) with a resistance of 4-7 MΩ. The pipettes (pulled using a DMZ puller) were filled with a solution containing (in mM): 110 KCl, 10 NaCl, 1 MgCl2, 1 EGTA, and 10 HEPES, pH adjusted to 7.3 with KOH. The extracellular solution contained (ECS, in mM): 140 NaCl, 4 KCl, 2 CaCl2, 1 MgCl2, 4 Glucose, and 10 HEPES, pH adjusted to 7.4 with NaOH. To elicit action potentials, neurons were recorded in current-clamp configuration while being subjected to a series of incremental current injections. Each sweep began with a 50 ms baseline at 0 pA, followed by a hyperpolarizing pulse of −100 pA for 100 ms. Subsequently, depolarizing current steps were applied in 50 pA increments (ranging from 0 to 950 pA) for 100 ms, after which the holding current was returned to 0 pA.

For indentation experiments, neuronal cell membranes were displaced ten times within a range of 1 – 9 µm (empirically measured membrane displacements were 0.6, 1.6, 2.6, 4, 4.6, 5.6, 6.6, 7.2, 8, 9 µm) using a heat-polished borosilicate glass pipette (mechanical stimulator) driven by an MM3A micromanipulator (Kleindiek Nanotechnik, Germany)^32^. All dishes were incubated in 1:400 Isolectin-GS-IB4-AlexaFluor 594 conjugate (I21413, Thermo Fischer Scientific) for 10 min followed by a 3 min wash in ECS. Labelled IB4^+^ neurons were identified under a Zeiss 200 upright epifluorescence microscope (40x objective) connected to a CoolSnapEZ camera (Photometrics). For a neuron to be considered “responder” to mechanical indentation, it had to respond to the all-subsequent force stimuli after its first response to an indentation. A “non-responder” neuron did not show currents between the 1-9 um of indentation tested. The first force at which the neurons responded was their mechanical threshold. Mechanically evoked currents were further characterized according to time constant to inactivation into rapidly adapting (RA, T_inact_ < 10 ms), intermediately adapting (IA, T_inact_ > 10 < 50 ms) or slowly adapting (SA, T_inact_ > 50 ms) currents.

Neurons were first tested for action potential properties, then switched to voltage-clamp mode for indentation experiments. Current-clamp and voltage-clamp traces were analyzed using FitMaster (HEKA, Elektronik GmbH, Germany).

### siRNA transfection experimental design

Two hours after plating the DRG neurons in glass-bottomed dishes, siRNA transfection was carried out using DharmaFECT (Horizon) reagents according to the manufacturers’ guidelines. Briefly, siRNAs were resuspended in serum- and antibiotic-free DRG neuron plating medium in a tube (total volume of 100 µL per dish), and 4.5 µL of DharmaFECT 4 transfection reagent was mixed with serum- and antibiotic-free DRG neuron plating medium (total volume of 100 µL) in another tube. Each tube was incubated separately for 5 min at RT and then mixed and incubated for an additional 20 min. Antibiotic-free complete medium (800 µL) was added to the transfection mix before being added to the neurons. Experimental dishes were transfected with a final concentration of 50 nM ON-TARGETplus SMARTpool mouse *B3gnt2* or mouse *Rrad* and 50 nM of siGLO green transfection indicator, and control dishes were transfected with 50 nM of ON-TARGETplus Non-targeting Pool and 50 nM of siGLO green transfection indicator. Transfected neurons that were not live stained for IB4 were patched 48 hours after transfection with siRNA.

### Immunocytochemistry

Cultured DRG neurons in PPS membrane slides were washed once for 5 min with PBS-Tween and fixed with Zamboni’s fixative for 10 minutes at room temperature (∼22 deg C). Post-fixation, slides were washed three times for 5 minutes with PBS-Tween, permeabilized with 0.05% Triton X-100 for 5 min and washed again three times for 5 minutes each. Before staining, slides were blocked in antibody diluent solution: 0.2% (v/v) Triton X-100, 5% (v/v) donkey serum and 1% (v/v) bovine serum albumin in PBS for 1 hour at room temperature before overnight incubation at 4 °C with 1:500 polyclonal anti-goat TrkA (R&D systems, AF1056) primary antibody. The next day, slides were washed three times using PBS-Tween and incubated with anti-goat conjugated secondary antibody (1:500 Alexa Fluor 488, Jackson Immuno Research Labs 705545147) and 1:200 Isolectin-GS-IB4-AlexaFluor 594 conjugate (I21413, Thermo Fischer Scientific) for 2 hours and then washed three times with PBS-Tween. Finally, slides were washed once with distilled water, air dried, and stored at 4 °C until laser microdissection (LMD).

### Immunohistochemistry of sections

L3 and L4 DRG were collected from male mice, postfixed in Zamboni’s fixative for 1 hour, and incubated overnight in 30% (w/v) sucrose (in PBS) at 4°C for cryoprotection. DRG were next embedded in Shandon M-1 Embedding Matrix (Thermo Fisher Scientific), snap frozen on dry ice, and stored at −80 °C. Embedded DRG were sectioned (5 μm) using a Leica Cryostat (CM3000; Nussloch) and mounted on polyethylene naphthalate (PEN) glass frame membrane slides (Leica microsystems). Consecutive sections were collected on different slides to prevent sampling the same neuronal cell body multiple times. During staining, slides were washed with PBS-Tween, blocked in antibody diluent solution for 1 h, and then incubated with primary antibodies at 4 °C. The following primary antibodies were used: anti-P2X3 (host: guinea pig, Alomone Labs APR-016-GP, 1:200), anti-CGRP (host: rabbit, Immunostar 24112, 1:1000), anti-NF200 (host: chicken, Abcam AB72996, 1:1000). After washing the unbound primary antibodies as described for immunocytochemistry, slides were incubated for 2 h at room temperature with the following species-specific conjugated secondary antibodies at 1:500: Alexa Fluor 488 anti-rabbit (A21206), Alexa Fluor 568 anti-guinea pig (A11075), and Alexa Fluor 488 (A11039). The unbound secondary antibodies were then thoroughly washed with PBS-Tween three times, and then once with distilled water and air-dried. The sections were imaged using an Olympus CSU-WI spinning disk microscope at 20x magnification. Exposure levels were kept constant for each slide, and the same contrast enhancements were made for all slides. Stained and imaged slides were stored at 4 °C until LMD.

### Image segmentation

The images obtained from DRG tissue sections were imported to the QuPath software with an integrated Cellpose extension (as detailed in https://github.com/BIOP/qupath-extension-cellpose) for further analysis. We previously trained our Cellpose 2.0^46^ model (“trk_test”) based on the cyto2 model using a human-in-the-loop approach with a mean diameter of 30. This model was integrated into QuPath^47^ to segment neurons positive for specific antibody staining. The contours thus obtained were dilated radially to prevent accidental laser ablation and converted to an LMD-compatible format for automated LMD^17^ (as detailed in https://github.com/CosciaLab/Qupath_to_LMD).

### Laser microdissection

We collected cultured cells and tissue samples using the Leica LMD 7 system and Leica Laser Microdissection software (version 8.3.0.08259). Cell culture and tissue samples were cut using a 40x objective in fluorescence or brightfield mode. For tissue samples, we used the following laser settings: power 60, aperture 1, speed 26, head current 45%, pulse frequency 2110. For the cell culture samples, we used the following laser settings: power 60, aperture 1, speed 20, head current 37%, pulse frequency 1537. Low-binding 384-well plates ((Eppendorf, cat.no. 0030129547) were used for sample collection and configured over the “universal holder” function with one empty well between the samples. For pooled sample analysis we cut ∼100 neuron sections from tissue and ∼50 cultured neurons of each of the three subsets from each of the three mice. Different cohorts of mice were used for tissue and cell culture experiments. For single-neuron analysis, 18 tissue sections were taken from each of three mice in each of the three neuron subsets.

### Sample preparation for LC-MS analysis

Sample preparation was performed as previously described^17^. Briefly, before sample preparation, 15 μl of acetonitrile (HPLC grade, VWR, cat. 83640.290) was added to each well, vortexed, and dried using speedvac. This drying process ensured that the sample contours were concentrated at the bottom of each well. Pipetting was performed either manually or using the MANTIS Liquid Dispenser (Formulatrix, V3.3 ACC RFID, software version 4.7.5) and high-volume diaphragm chips (Formulatrix, cat.no. 233128 ).

For cell culture, samples were first incubated in 4 μl 60mM Triethylammonium bicarbonate pH 8.5 (TEAB; Merck, cat.no. T7408-100ML) at 95 °C for 60 min. Then, the samples were cooled, and 1μl of 100% ACN (HPLC-grade, VWR, cat. 83640.290) was added and incubated for 60 min at 75 °C. Next, 1 μl of LysC (Promega, cat. no VA1170) (2 ng/μL) was added, followed by a 2 h digestion at 37 °C. Then, 1 μl of trypsin (Promega, cat.no. V5117) (2 ng/µL) was added with overnight incubation at 37 °C in the thermal cycler. Digestion was stopped the next day by addition of trifluoroacetic acid (TFA, final concentration 1% v/v) (cat. no. 96924-250ML-F). The samples were vacuum dried before peptide clean-up.

For tissue samples, the lysis buffer for sample preparation consisted of 0.1% n-dodecyl-beta-maltoside (DDM; Sigma-Aldrich, cat.no. D4641-500MG), 5mM Tris (2-carboxyethyl) phosphine hydrochloride (TCEP; Sigma-Aldrich, cat.no. C4706-2G), 20mM 2-chloroacetamide (CAA; Sigma-Aldrich, cat.no. C0267-100G and 0.1M TEAB in water. 2μL of lysis buffer were added to wells containing samples and incubated for 60 min at 95 °C. Subsequently, 1 μL of LysC (Promega, cat.no VA1170) (2 ng/μL) was added and digested for 2 h at 37°C, followed by addition of 1 μl of trypsin (Promega, cat.no. V5117) (2 ng/ul) and overnight digestion at 37 °C in a thermal cycler. After digestion, the reaction was stopped by adding trifluoroacetic acid (TFA, final concentration 1% v/v) (cat. 96924-250ML-F). The samples were then vacuum-dried before peptide clean-up.

### Peptide clean-up with C-18 tips

Peptide clean-up was performed using the Evotip Pure protocol, as recommended by the manufacturer. Briefly, Evotips (EV2013, Evotip Pure, Evosep, cat.no. EV2013) were first washed with 20 μL of buffer B (99.9% ACN, 0.1% FA) and centrifuged at 700 rpm for 1 min. Then, 20 μL of buffer A (99.9% water, 0.1% FA) was added to each C-18 tip, activated in isopropanol for 20 s, and centrifuged again at 700 rpm for 1 min. Digested tissue or cell culture samples were then loaded onto Evotips, washed once with 20 μL buffer A, and finally eluted with 20 μl buffer B in a 96-well plate (Thermo Fisher Scientific, cat. no. AB1300), and vacuum dried (15 min at 60 °C). The samples were stored at 20 °C until liquid chromatography–mass spectrometry (LC–MS) analysis. For LC-MS analysis, 4.2 μL of MS loading buffer (3% acetonitrile, 0.1% TFA in water) was added, the plate was vortexed for 10 s, and centrifuged for 1 min at 700 g. Four microliters were injected into the mass spectrometer for LC-MS analysis.

### Liquid chromatography–mass spectrometry (LC – MS) analysis

LC-MS analysis was performed with an EASYnLC-1200 system (Thermo Fisher Scientific) connected to a trapped ion mobility spectrometry quadruple time-of-flight mass spectrometer (timsTOF SCP (cell culture samples), timsTOF Ultra (tissue samples), Bruker Daltonik) with a nano-electrospray ion source (CaptiveSpray, Bruker Daltonik). An autosampler was configured to pick samples from 96-well plates. Peptides were loaded onto a 20-cm home-packed HPLC column (75-mm inner diameter packed with 1.9-mm ReproSil-Pur C18-AQ silica beads, Dr. Maisch).

For tissue samples, peptides were separated using a linear gradient from to 7-30% buffer B (0.1% formic acid and 90% ACN in LC-MS grade water) for 14 min, followed by an increase to 60% for 1 min and a 1.5-minute wash in 90% buffer B at 250 nL/min. Buffer A consisted of 0.1% formic acid in liquid chromatography-mass spectrometry (LC-MS) grade water. The total gradient length was 21 min. A column oven was used to maintain the column temperature at 40 °C.

For cell culture analysis, peptides were separated using a linear gradient from to 4-30% buffer B (0.1% formic acid and 90% ACN in LC-MS grade water) for 47 min, followed by an increase to 60% for 3 min and a 3 min wash in 90% buffer B, followed by a gradual decrease to 50% buffer B over 7 min. The flow rate was 250 nL/min.

For tissue samples, we used a dia-PASEF method with 8 dia-PASEF scans separated into 3 ion mobility windows per scan covering a 400-1000 m/z range by 25 Th windows and an ion mobility range from 0.64 to 1.37 Vs cm^-2^. The mass spectrometer was operated in high sensitivity mode, with an accumulation and ramp time at 100 ms, capillary voltage set to 1750V and the collision energy as a linear ramp from 20 eV at 1/k_0_ = 0.6 Vs cm^-2^ to 59 eV at 1/k_0_ = 1.6 Vs cm^-2^.

For cell culture analysis, we used a dia-PASEF long method with 16 dia-PASEF scans separated into 2 ion mobility windows per scan covering a 400-1000 m/z range by 26 Th windows and an ion mobility window range from 0.60 to 1.60 cm^-2^. The mass spectrometer was operated in high sensitivity mode, with an accumulation and ramp time at 100 ms, capillary voltage set to 1600V and the collision energy as a linear ramp from 20 eV at 1/k_0_ = 0.6 Vs cm^-2^ to 59 eV at 1/k_0_ = 1.6 Vs cm^-2^.

### Mass spectrometry raw file analysis

We used DIA-NN^6^ (version 1.8.1) for the dia-PASEF raw file analysis and spectral library generation. For spectral library generation to analyze dia-PASEF data, the mouse FASTA file was downloaded from Uniprot (2023 release, UP000000589_10090 Mus Musculus, downloaded on August 2nd). DIA-NN in silico predicted libraries were generated using the mouse FASTA file, and frequently found contaminants^62^ (mouse + mouse tissue contaminants). Deep learning-based spectra, RTs, and IMs prediction were enabled for the appropriate mass range of 300-1000 m/z. N-terminal M excision and cysteine carbamidomethylation were enabled as fixed modifications. A maximum of two missed cleavages were allowed, and the precursor charge was set to 2–4. The library consisted of 22092 protein isoforms, 34222 protein groups, and 6776851 precursors in 3121869 elution groups.

DIA-NN^63^ was operated in the default mode with minor adjustments. Briefly, MS1 and MS2 accuracies were set to 15.0, scan windows to 0 (assignment by DIA-NN), isotopologues were enabled, no MBR for project-specific DIA-NN-refined libraries, heuristic protein inference, and no shared spectra. Proteins were inferred from genes, neural network classifier was set to single-pass mode, quantification strategy as ‘Robust LC (high precision).’ Cross-run normalization was set to ‘RT-dependent,’ library generation as ‘smart profiling,’ speed and RAM usage as ‘optimal results.’

### Quantification and statistical analysis

Proteomics data analysis was performed with Perseus^64^ (version 1.6.15.0) and within the R environment (https://www.r-project.org/) with the following packages: tidyverse (version 2.0.0), UpSetR (version 1.4.0), FactoMineR (version 2.11), factoextra (version 1.0.7), ComplexHeatmap (version 2.18.0), RColorBrewer (version 1.1.3), circlize (version 0.4.16), dendsort (version 0.3.4), ggrepel (version 0.9.5), viridis (version 0.6.5) and clusterProfiler (version 4.6.2). Before statistical analysis, data were filtered to retain only proteins with 70% non-missing data in at least one group (if not stated otherwise in the figure legend). Missing values were imputed based on a normal distribution (width = 0.3, downshift = 1.8) before statistical testing. For single-neuron proteomics (Fig. 3), two outlier samples (ids .6944 and .7041) were removed and samples only kept for further analysis with a minimum of 700 protein groups, resulting in 20 CGRP^+^ peptidergic and 22 P2X3^+^ non peptidergic neuron proteomes. For multiple group comparisons, analysis of variance (ANOVA) was used. For both tests, a permutation-based FDR of 5% was applied to correct multiple hypothesis testing. Pathway enrichment analysis was performed in Perseus based on Fisher’s exact test (categorical data) or 1D pathway enrichment analysis^65^. Hallmark gene sets, WikiPathways, and Reactome pathways were enriched terms filtered using a Benjamini-Hochberg FDR cut-off of 0.05. The minimum category size was set to five.

To estimate transcriptome vs proteome depth in Extended Data Fig 3, we downloaded the scRNA seq GEO dataset from Sharma et al (GSM4130750_WT_1.csv) ^2^. CGRP and Non-peptidergic neurons were then grouped to calculate the average transcript reads from all genes in each subset. These genes were then matched by name to our proteomics dataset. Data were plotted as a histogram across the average transcript reads showing the proportion of gene counts detected in proteomics over transcriptomics. Furthermore, we selected some genes with low (<100 transcripts) and high (>1000 transcripts) average copy numbers to demonstrate the robust detection of proteins from genes with low transcript reads.

For proteomic-transcriptomic comparisons in Fig 2, we used data from Bhuiyan et al^3^. Briefly, the “Marker Genes” file was downloaded from the accompanying website (https://painseq.shinyapps.io/harmonized_painseq_v1/) and DRG neuron and non-neuron tabs were used for further analysis in R or Perseus. To match non-neuronal gene names between transcriptomics and proteomics data, we compiled all non-neuronal associated genes from all types of DRG associated non-neuronal cells (Fig 2e). These were matched with our proteome dataset to calculate the number of non-neuronal proteins. The mean values of the matched proteins were plotted in a sequential color map.

To compare neuronal marker genes (Fig 2f), we compiled a list of the top 50 transcriptomics-related genes from all 18 subsets of sensory neurons (with all positive log fold change values), matched them against our proteomics data, and performed z-scoring based on spearman distance for hierarchical clustering. For single CGRP^+^ neuron comparisons, all genes with a positive log fold change value (obtained from the Marker Genes file in Bhuiyan et al) in each of the CGRP-related transcriptomic cluster were normalized between 0 and 1 using the preProcess() function of the caret library in R using the range method. Similar normalization was carried out for the log2 intensity of proteins quantified in each of the CGRP^+^ single neurons. The names of the genes were matched to get a normalized value from transcriptomics (Gtn) and a normalized value from proteomics (Gpn) for each matching gene. For each transcriptomics cluster we then calculated a Euclidean distance score 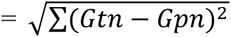 (resulting in a 20×6 matrix) to estimate the probability that our single neuron belongs to a specific transcriptomics defined CGRP cluster and visualized using a heatmap.

Statistical tests for electrophysiological data are detailed in the figure legends. Briefly, comparisons between two groups were carried out using Student’s t-test (with Welch’s correction if the variances were significantly different between groups), and proportions were compared using the chi-sq test in GraphPad Prism 9.

## Data availability

The mass spectrometry proteomics data have been deposited to the ProteomeXchange Consortium via the PRIDE partner repository^66^ with the dataset identifier PXD070495. All other data will be shared upon request by the corresponding authors.

## Code availability

The code used for analysis and plots were adapted from standard R packages as detailed in the Methods section. More detailed information is available upon request.

## Acknowledgements

This work was funded by an EFIC-Grünenthal award to S.C., an ERC senior award “Grant #101142488, “Pain Channels” to GRL. FC acknowledges funding support by the Federal Ministry of Education and Research (BMBF), as part of the National Research Initiatives for Mass Spectrometry in Systems Medicine (grant agreement No. 161L0222) and funding from the European Research Council (ERC) under the European Union’s Horizon 2020 research and innovation program (grant agreement No. 101115681).

## Author contributions

Conceptualization: S.C., F.C., G.R.L.

Cell culture, path clamp electrophysiology, immunocytochemistry, image analysis: S.C.

Tissue processing, immunohistochemistry and laser microdissection: S.C. with help from A.M., W.L., L.W.

Proteomics sample preparation, processing and analysis: A.M. with help from F.C. and S.C. Writing: S.C., F.C. and G.R.L. with help from all authors

Funding: S.C., F.C., G.R.L.

**S.C., G.R.L. and F.C. are inventors in an EU patent filed based on this work. The authors declare no other conflicts of interest.**

**Table Legends:**

Extended Data Table 1: Top 50 marker proteins based on subset-specific sensory neuron proteomics.

Supplementary Data Table 1: List of transmembrane proteins and ion channels identified in different subsets of sensory neurons.

Supplementary Data Table 2: Z-scored list of ANOVA significant proteins from sensory neuronal subsets.

Supplementary Data Table 3: Unimputed list of proteins identified from intact, pooled peptidergic and non-peptidergic nociceptors.

Supplementary Data Table 4: Unimputed list of proteins identified from pooled and single intact mechanoreceptors.

Supplementary Data Table 5: Unimputed list of proteins identified in single sections of nociceptors.

Supplementary Data Table 6: List of upregulated proteins after inflammation in peptidergic, non-peptidergic nociceptors and mechanoreceptors.

## Extended Data Figures

**Extended Data Fig 1:**
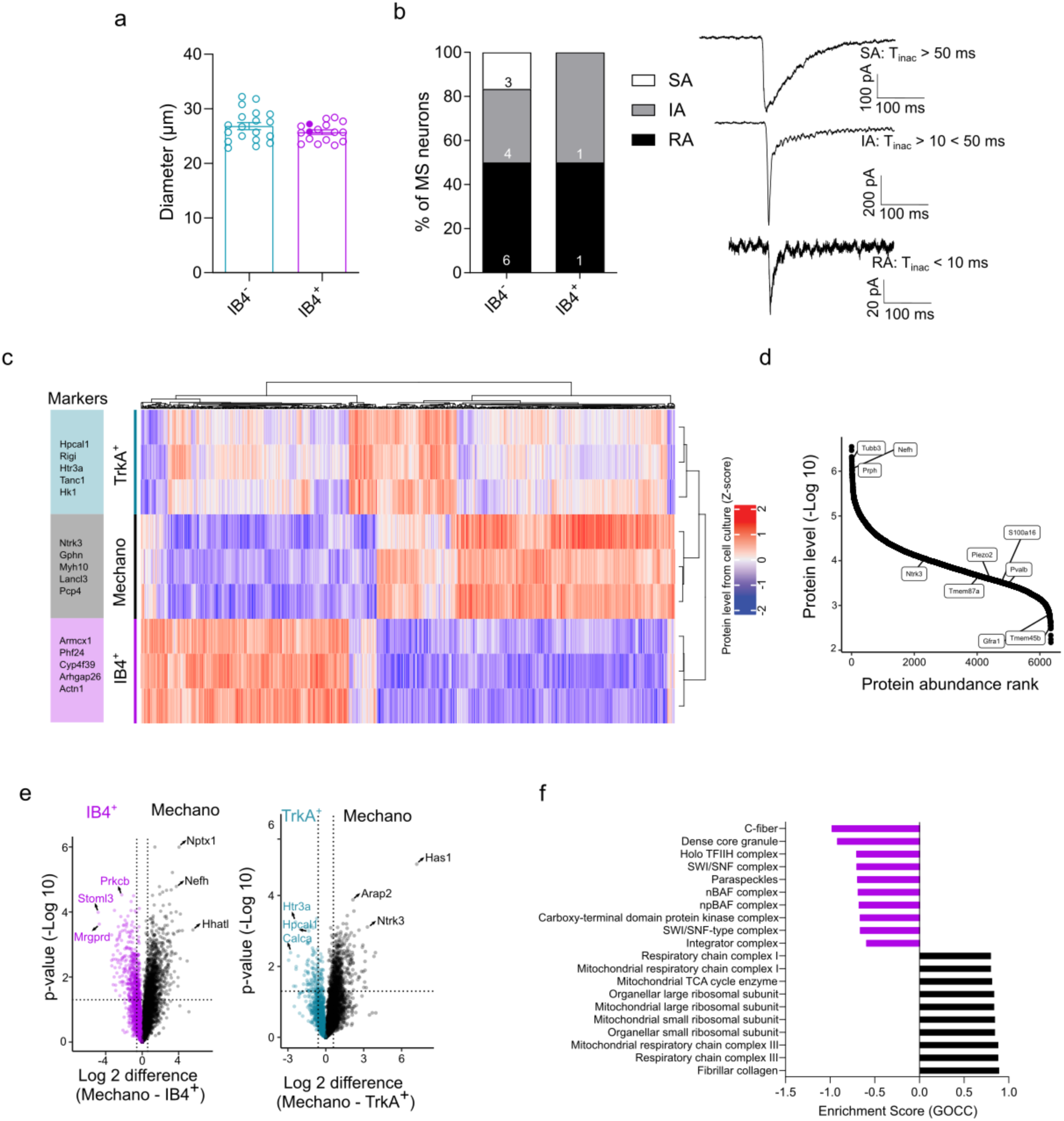
Proteomics-based, subset-specific markers of sensory neurons. **a)** Bar plot showing the distribution of the diameters of patched nociceptors. Filled dots in IB4+ neurons represent the two mechanosensitive non-peptidergic neurons. **b)** Contingency plot of percent mechanically active currents with slowly adapting (SA), intermediately adapting (IA) and rapidly adapting (RA) biophysical properties along with their representative traces. **c)** Hierarchical clustering of proteins identified in cultured peptidergic and non-peptidergic nociceptors and mechanoreceptors. The top five proteins in each cluster are highlighted on the left. Each column represents z-scored protein levels. **d)** Dynamic range of protein abundance in the mechanoreceptor subset. **e)** Volcano plot showing the pairwise proteomic comparison between non-peptidergic (left) or peptidergic (right) nociceptors and mechanoreceptors with marker proteins highlighted. **f)** Pathway enrichment analysis based on t-test differences between non-peptidergic nociceptors and mechanoreceptors. The top ten pathways with a Benjamini-Hochberg false discovery rate (FDR) < 0.05 are shown. Cyan = peptidergic, magenta = non-peptidergic, black = mechanoreceptor. Error bars represent s.e.m.

**Extended Data Fig 2:**
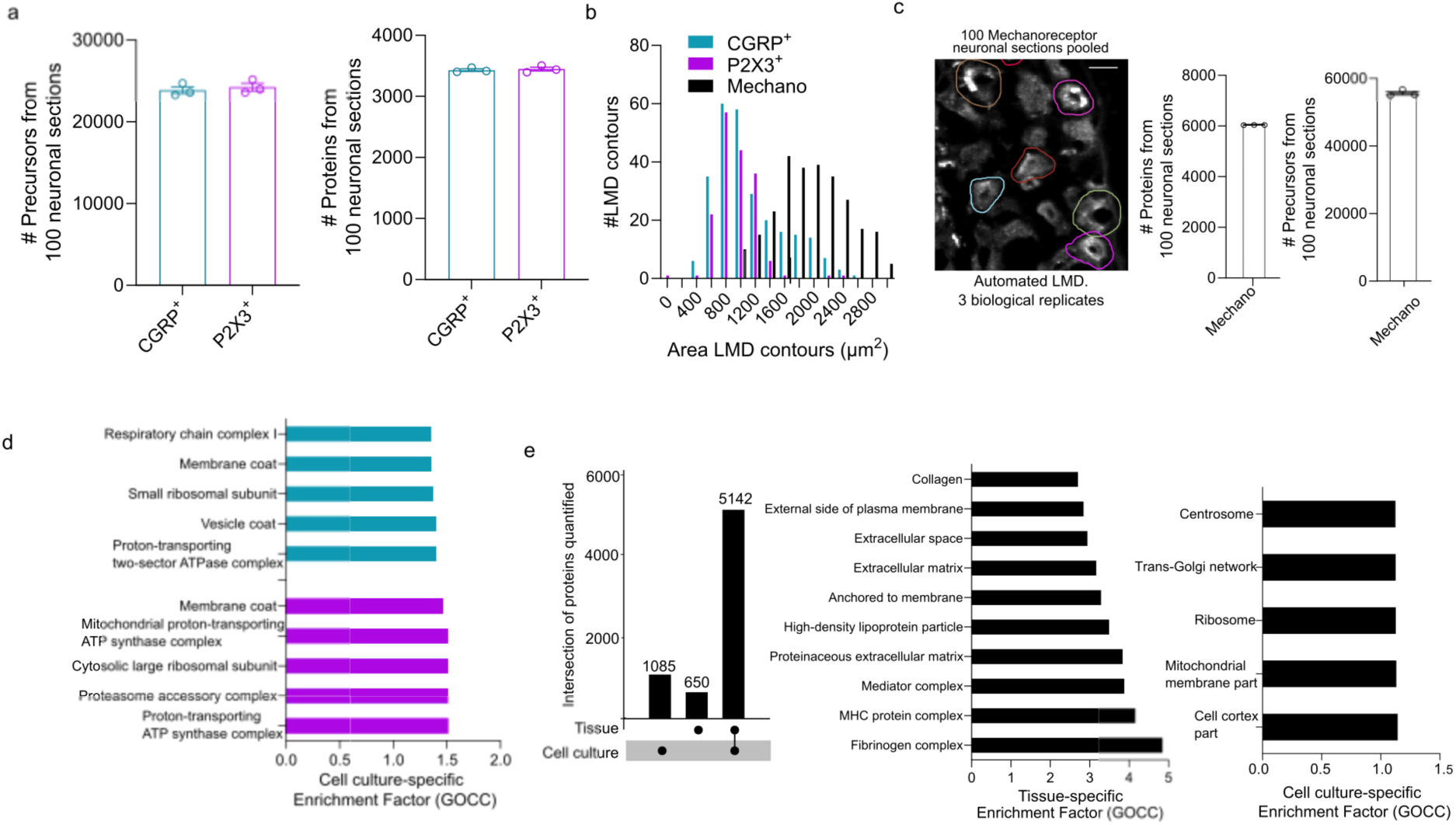
Protein quantification from intact sensory neuron tissue sections. **a)** Number of precursors and proteins identified from peptidergic (P2X3^−^/CGRP^+^) and non-peptidergic (P2X3^+^/CGRP ^−^) nociceptors. **b)** Histogram of area of the contours isolated by automated laser microdissection (LMD) from nociceptors and mechanoreceptors. **c)** Representative image of NF200 stained dorsal root ganglion (DRG) tissue sections with highlighted LMD contours (left). Number of precursors and proteins identified from the mechanoreceptors (right). **d)** Pathway enrichment analysis (Fischer’s exact test) of uniquely identified proteins from cultured nociceptors. **e)** Upset plot showing common and unique proteins quantified from mechanoreceptors in intact tissue and in culture (left), along with pathway enrichment analysis (Fischer’s exact test) of uniquely identified proteins from intact, tissue-derived (middle) and cultured (right) mechanoreceptors. Error bars represent s.e.m.

**Extended Data Fig 3:**
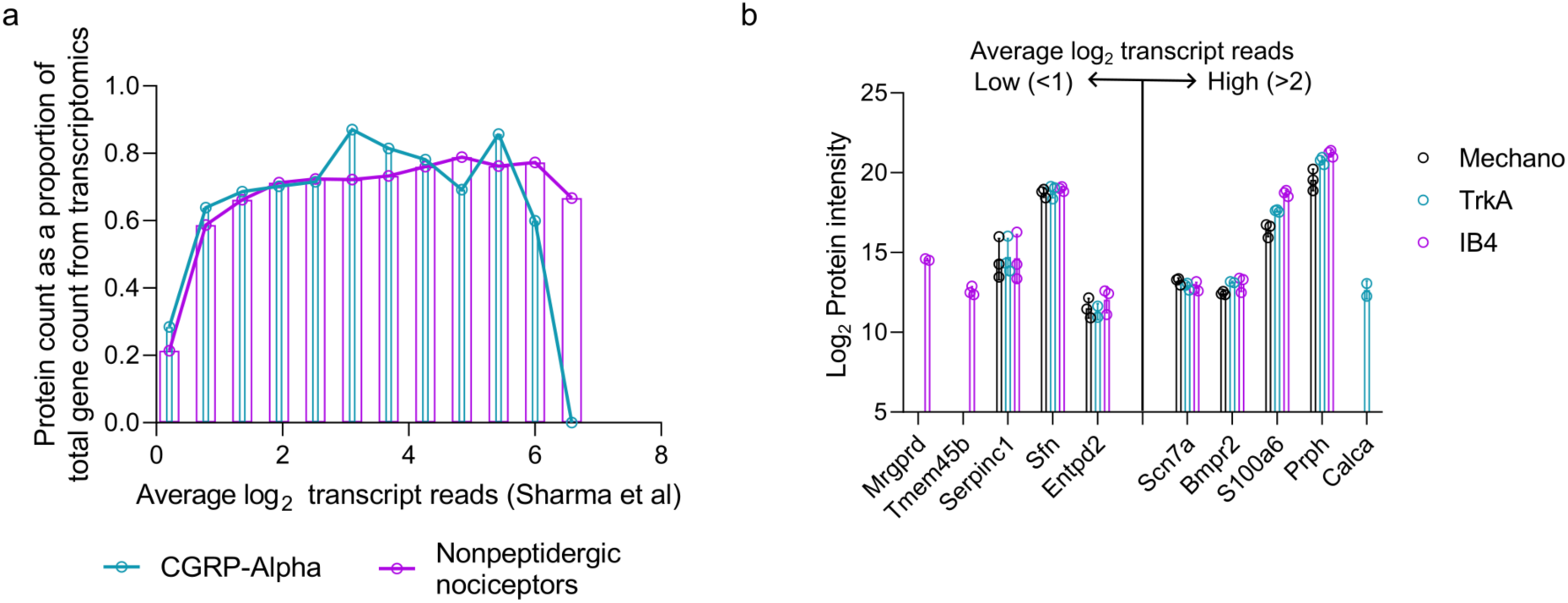
**Comparison of gene detection by transcriptomics and by proteomics**. **a)** Histogram showing genes detected in proteomics as a fraction of genes detected using transcriptomics (from Sharma et al’s scRNA seq dataset) across a range of log transformed average transcript reads. **b)** Plots of protein intensity for selected genes with low and high average transcript reads. Cyan = peptidergic, magenta = non-peptidergic, black = mechanoreceptor.

**Extended Data Fig 4:**
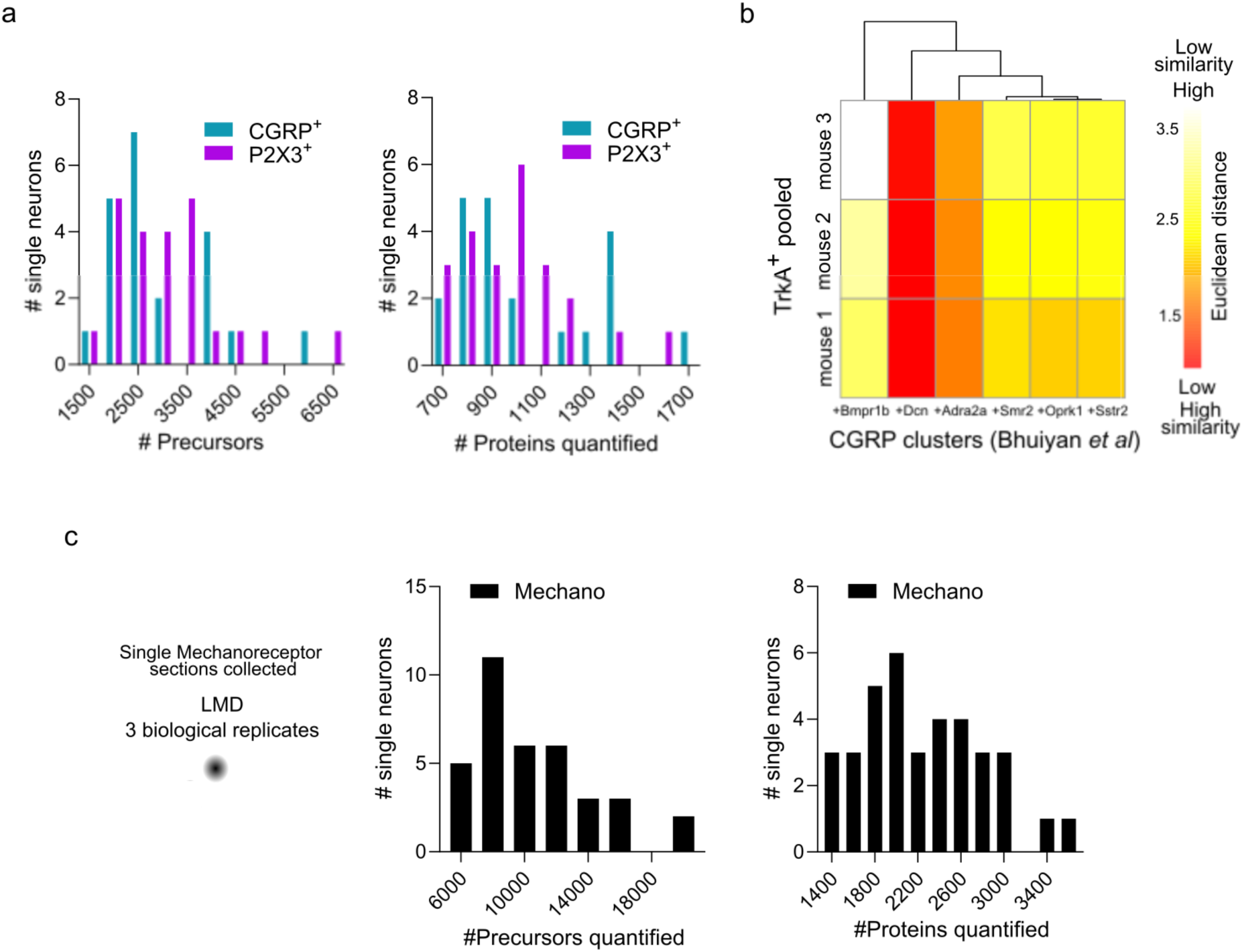
Protein quantification from single sensory neuron sections. **a)** Histograms showing the number of precursors and protiens quantified from each nociceptor section. **b)** Similarity plot based on Euclidean distance between enriched genes in CGRP (peptidergic) subclusters (columns) identified by single-cell transcriptomics, and proteome of pooled, cultured TrkA^+^ peptidergic nociceptors (row). Red = high similarity; white = low similarity. **c)** Histogram showing the number of precursors and proteins quantified from each individual mechanoreceptor section.

**Extended Data Fig 5:**
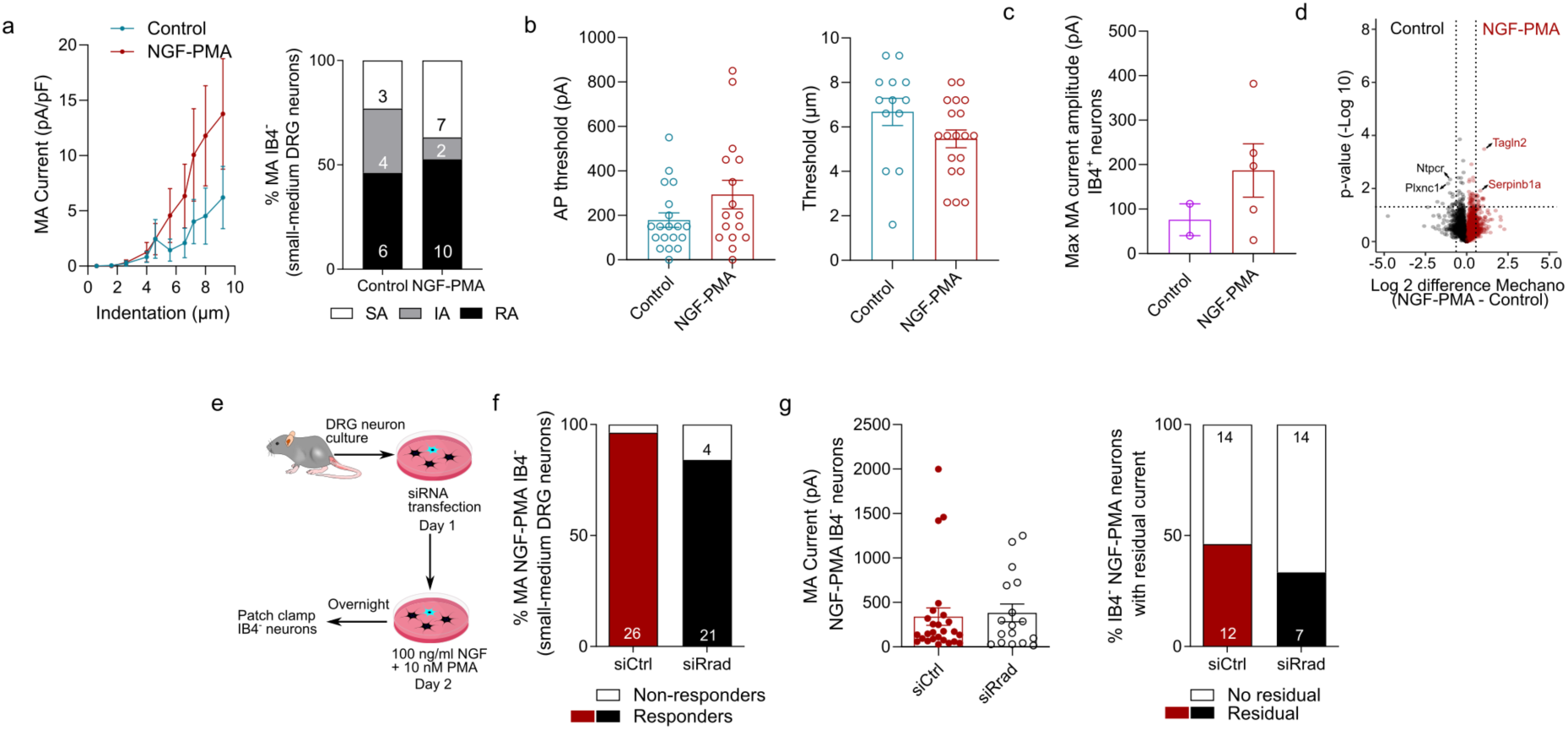
Rrad does not ameliorate inflammation-induced mechanical hypersensitivity of peptidergic nociceptors. **a)** Mechanically active (MA) current amplitude at across a range of indentation in control and NGF-PMA sensitized peptidergic nociceptors, along with quantification of the proportion of slowly adapting (SA), intermediately adapting (IA) or rapidly adapting (RA) currents elicited. **b)** Action potential generation and mechanical indentation threshold of control (cyan) and NGF-PMA treated (red) peptidergic nociceptors. **c)** Bar graph showing indentation-evoked mechanosensitive current amplitudes of control (magenta) and inflamed (red) non-peptidergic nociceptors. **d)** Volcano plot showing the pairwise proteomic comparison between control and NGF-PMA treated mechanoreceptors with the top regulated proteins highlighted. **e)** Schematic workflow of the validation experiment by knockdown of Rrad using siRNA prior to NGF-PMA treatment and patch-clamp electrophysiology of peptidergic neurons. **f)** Percentage of mechanically sensitive NGF-PMA treated peptidergic nociceptors transfected with control siRNA (red) and Rrad-targeting siRNA (black). The number of neurons is indicated in the bars. **g)** Current amplitudes (left) and percentage of neurons with residual currents in NGF-PMA treated peptidergic nociceptors transfected with control siRNA (red) and Rrad-targeting siRNA (black). Error bars represent s.e.m.

